# The DNA Damage Response kinase ATM restricts Golgi extension

**DOI:** 10.64898/2026.03.23.713647

**Authors:** Caroline Soulet, Julie Catalan, María Moriel-Carretero

## Abstract

The master kinases of the DNA damage response (DDR), ATR, ATM and DNA-PK, become active in response to DNA damage and orchestrate a downstream wave of phosphorylations contributing to DNA damage repair and preservation of cellular homeostasis. Of them, we recently demonstrated that ATM binds the pool of the lipid phosphatidyl-inositol-4-phosphate (PI4P) situated at the Golgi membrane. Depending on PI4P availability at Golgi membranes, ATM is more or less titrated away from the nucleus, which translates into responses to nuclear DNA damage of matching intensity. Building on this knowledge, in this work we asked if, beyond the Golgi merely serving as a docking platform that retains ATM away from the nucleus, ATM does exert any role important for Golgi biology. We found that ATM maintains Golgi morphology by counteracting its excessive deployment. This occurs both by its mere presence (likely antagonizing the Golgi-stretching action of the protein GOLPH3) and by phosphorylating Golgi-resident substrates. Of relevance, we also report that the morphological alterations caused to the Golgi without ATM affect the biology of a model Golgi cargo. Our findings nourish the growing evidence that kinases of ATM’s family display functional interactions with membranes and highlights an underappreciated crosstalk between the Golgi and the nucleus.

## Introduction

The master kinases of the DNA damage response (DDR), ATR (ataxia telangiectasia and Rad3-related kinase), ATM (ataxia telangiectasia-mutated kinase) and DNA-PK (DNA-dependent protein kinase) become active in response to DNA damage and orchestrate a downstream wave of phosphorylations contributing to DNA damage repair and preservation of cellular homeostasis (1). They belong to the family of phosphatidyl-inositol-3-kinase-related kinases (PIKKs), yet no longer display the ability to phosphorylate phosphatidylinositol (PI) moieties (2). However, it is formally possible that they have kept their ability to interact with lipid moieties. In this sense, the PIKK kinase mTOR indeed binds to lipids within the lysosome membrane, a step that is key for its proper activation (3). ATR connection to membranes has also been reported in scattered works: its yeast counterpart Mec1 binds and phosphorylates targets on the outer mitochondrial membrane upon nutrient shortage (4); ATR localizes to and becomes active at the NE upon nuclear swelling caused by hyper-osmotic stress (5); responds to an increase in membrane fluidity upon excess of polyunsaturated fatty acids (6); and monitors accurate membrane scission at the cytokinetic site (7). Further, we showed that ATR’s activity is influenced by the type and quantity of phospholipids in the nuclear space (8). Regarding the DDR kinase ATM, we recently demonstrated that ATM binds to the pool of the lipid phosphatidyl-inositol-4-phosphate (PI4P) situated at the Golgi membrane. Depending on PI4P availability at Golgi membranes, ATM is more or less titrated from the nucleus, which translates into either weaker or more robust responses to nuclear DNA damage, respectively (9). It is important to highlight that the Golgi is a very dynamic organelle with a constantly changing and extreme plasticity. Together, a picture emerges in which PIKKs can, not only interact with membranes *via* their potential lipid-binding abilities, but perhaps probe and be sensitive to mechanical properties of the membranes harboring those lipids.

In this work, we built on our discovery that a pool of ATM is Golgi-resident to ask if, beyond the Golgi merely serving as a docking platform onto which to be retained away from the nucleus, ATM fulfils any role at the Golgi that could impact its biology. We found that ATM absence entails a deployment of this organelle dependent on the protein GOLPH3. GOLPH3 (also referred to in the literature as Vps74, GPP34, MIDAS, or GMx33α) is a robust PI4P binder at the Golgi, an interaction thanks to which it stretches this organelle by applying a strong tensile force (10). This raises the possibility that ATM absence permits GOLPH3 to better access PI4P. Beyond, we discover that ATM phosphorylates Golgi substrates to counteract excessive Golgi deployment. This “kinase mode” of control is modest under basal conditions and can become more marked upon cues instructing Golgi extension. Last, we find that the morphological alterations caused to the Golgi in the absence of ATM affect the biology of one model Golgi cargo. Our data provide yet another evidence that PIKKs display functional interactions with membranes, raise the question of how they sense mechanical properties of such membranes, and highlight (again) an undeniable crosstalk between different organelles, such as the Golgi and the nucleus.

## Results

### 1. ATM limits Golgi stretching, in part by counteracting GOLPH3

We have reported that the apical kinase of the DDR ATM can reside at the Golgi apparatus surface by virtue of its docking onto the resident lipid PI4P (9). This indirectly exerts regulatory effects on ATM’s nuclear duties, for it conditions its nuclear availability. Beyond this impact, we found it legitimate to ask if ATM fulfils any role at the Golgi that is important for this organelle. To assess this, we silenced ATM by siRNA in RPE-1 cells (Figure S1A), which did not lead to any detectable decrease in PI4P profiles at the Golgi, as estimated by immunofluorescence (Figure S1B). We measured the perimeter and the circularity of Golgi signals stemming from the *cis* and the *trans* Golgi faces (GM130 and TGN46, respectively). This is because, for a similar area, more or less deployed objects can be distinguished by their perimeter (see Annex 1 for explanation and quantification strategy). For the same reason, these deployed objects are prone to display a loss of circularity.

We observed a simultaneous increase in the perimeter of GM130 signals and a loss of circularity whenever ATM was absent (Figure 1A & B) in a reproducible manner (Figure 1C; and additional images in Figure S1C), and also for TGN46 signals (Figure S1C). This indicated that, in the absence of ATM, the Golgi becomes deployed. This was not due to ATM silencing altering cell cycle profiles, which could be postulated to make cells display different Golgi morphologies (Figure S1D).

**FIGURE 1.**
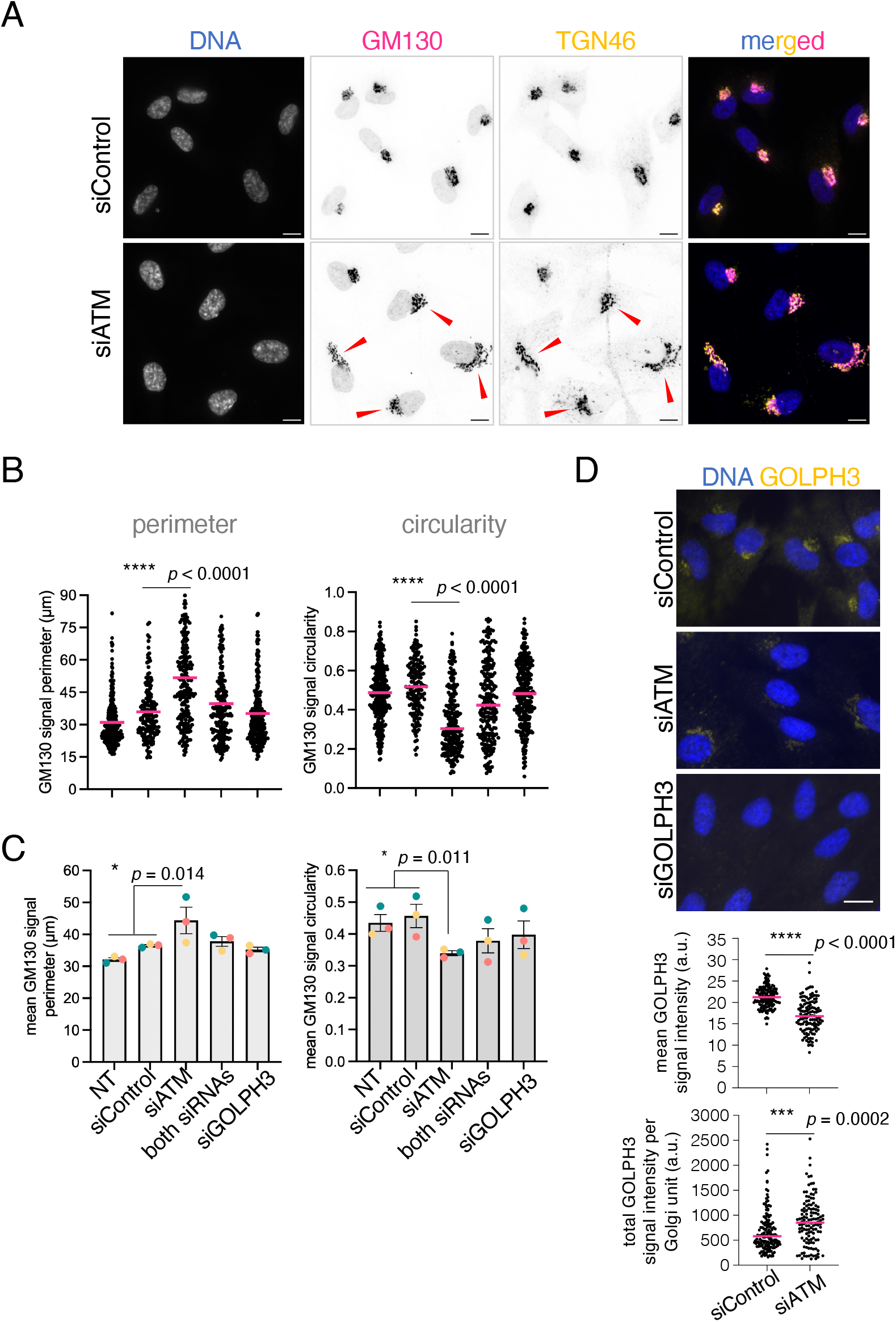
THE Golgi becomes deployed in the absence of atm. **A**.Immunofluorescence of the *cis*-Golgi marker GM130 and the *trans* Golgi marker TGN46 in cells transfected either with siControl or siATM. Bar size is 5 µm. Red arrowheads point to deployed Golgi units. **B**.The individual values for the perimeter (left) and the circularity (right) of individual Golgi unit signals (each point corresponds to one cell) as measured using GM130 signals from one experiment were plotted for the indicated conditions (where NT = non transfected). The horizontal pink bar indicates the median of the population. The statistics result from applying a non-parametric Mann-Whitney test comparing the medians of the indicated samples, since normality could not be guaranteed in all cases. **C**.The means from three independent experiments (three different color dots) as the one presented in (B) were used to assess the reproducibility of the phenotypes for the indicated conditions. The bar height indicates the mean of the means and the error bars reflect the standard error of the mean (SEM). The statistics reveal the outcome of an unpaired *t*-test comparing the mean of all control values against those obtained for siATM. **D**.GOLPH3 signals at Golgi units were measured using immunofluorescence in the indicated siRNA-treated cells. **Top**: images of representative cells. Size bar is 15 µm. **Bottom:** quantification of the mean GOLPH3 signal intensity per pixel and of total GOLPH3 signal intensity in each complete Golgi unit out of one individual experiment. Each dot corresponds to a Golgi unit from an individual cell. The horizontal pink bar indicates the median of the population. The statistics result from applying a non-parametric Mann-Whitney test comparing such medians.

Golgi deployment can be implemented by the protein GOLPH3 that, by binding to PI4P (as ATM), connects the *trans* Golgi with the actin cytoskeleton, thus mediating Golgi stretching (10). To ask if the Golgi extension seen in the absence of ATM depends on GOLPH3, we co-silenced both proteins (Figure S1A). We observed a partial reversion of the siATM-imposed Golgi extension, as illustrated by decreased Golgi perimeters (Figure 1B,C; Figure S1C), although the loss of circularity was not palliated. This suggested that, in the absence of ATM, GOLPH3 either gains more access to PI4P, or it is more active in its ability to extend the Golgi, or both. Also, the data imply that Golgi deployment in the absence of ATM is partly implemented by additional proteins beyond GOLPH3.

To challenge the notion that GOLPH3 may gain more access to the Golgi in the absence of ATM, we compared the GOLPH3 signals stemming from Golgi units in cells lacking (or not) ATM. The images shown in Figure 1D were not contrast-adjusted for GOLPH3 signals and were acquired with identical set-ups. The cells depleted for GOLPH3, used as a control, did not display any major signal either at the Golgi or the cytoplasm, in marked contrast with siControl cells, which displayed signals in both compartments (Figure 1D, top). In siATM cells, beyond clearly deployed Golgi units, GOLPH3 signals were markedly absent from the cytoplasm and neatly detectable at Golgi units (Figure 1D, top). Quantifications showed that GOLPH3 mean intensity per pixel decreased in siATM cells with respect to control ones (Figure 1D, bottom, left), which is an expected consequence of the deployment of the organelle. If the total GOLPH3 loaded onto each Golgi unit was however unchanged, we would expect an equal total intensity. However, the total GOLPH3 signal mapped on full Golgi units was increased in the absence of ATM (Figure 1D, bottom, right), and this in a very reproducible manner (Figure S1E), indicative of a global higher presence of GOLPH3. Thus, the absence of ATM allows GOLPH3 to better access Golgi units, which in part is responsible for an unscheduled Golgi extension.

### 2. ATM limits Golgi deployment through kinase-dependent and -independent modes

The effects seen in the absence of ATM appear to relate to this rendering available the PI4P moieties ATM would have bound, but they could be additionally due to the loss of ATM’s kinase activity. To clarify this, we inhibited ATM using the very specific molecule AZD0156, which abolishes its kinase activity but, by definition, does not get rid of the protein (as validated by western blot (Figure S2A, left, Ø)). By monitoring TGN46 signal morphology, we observed that ATM inhibition led to a neat deployment of the Golgi (Figure 2A) that was reproducible and of statistical significance (Figure 2B). Of note, inhibition of the other master DDR kinase, ATR, using the specific inhibitor VE-821, did not provoke any reproducible Golgi extension (Figure 2C). The deployment of Golgi units upon ATM inhibition (without any major impact on total ATM levels (Figure S2A, right, Ø)) was consistently observed in other cell types (Huh-7, of liver cancer origin; and A549, of lung cancer origin, Figure S2B), suggestive of a general effect. These data imply that ATM kinase activity is important under basal conditions to restrict Golgi extension, beyond the physical presence of ATM, which is an additional contributor basally limiting Golgi extension (the difference in mean perimeter is 3 μm between ATM-inhibited and untreated cells but of 7 μm between ATM-depleted and ATM-positive cells (Figure 1C, 2C)).

**Figure 2.**
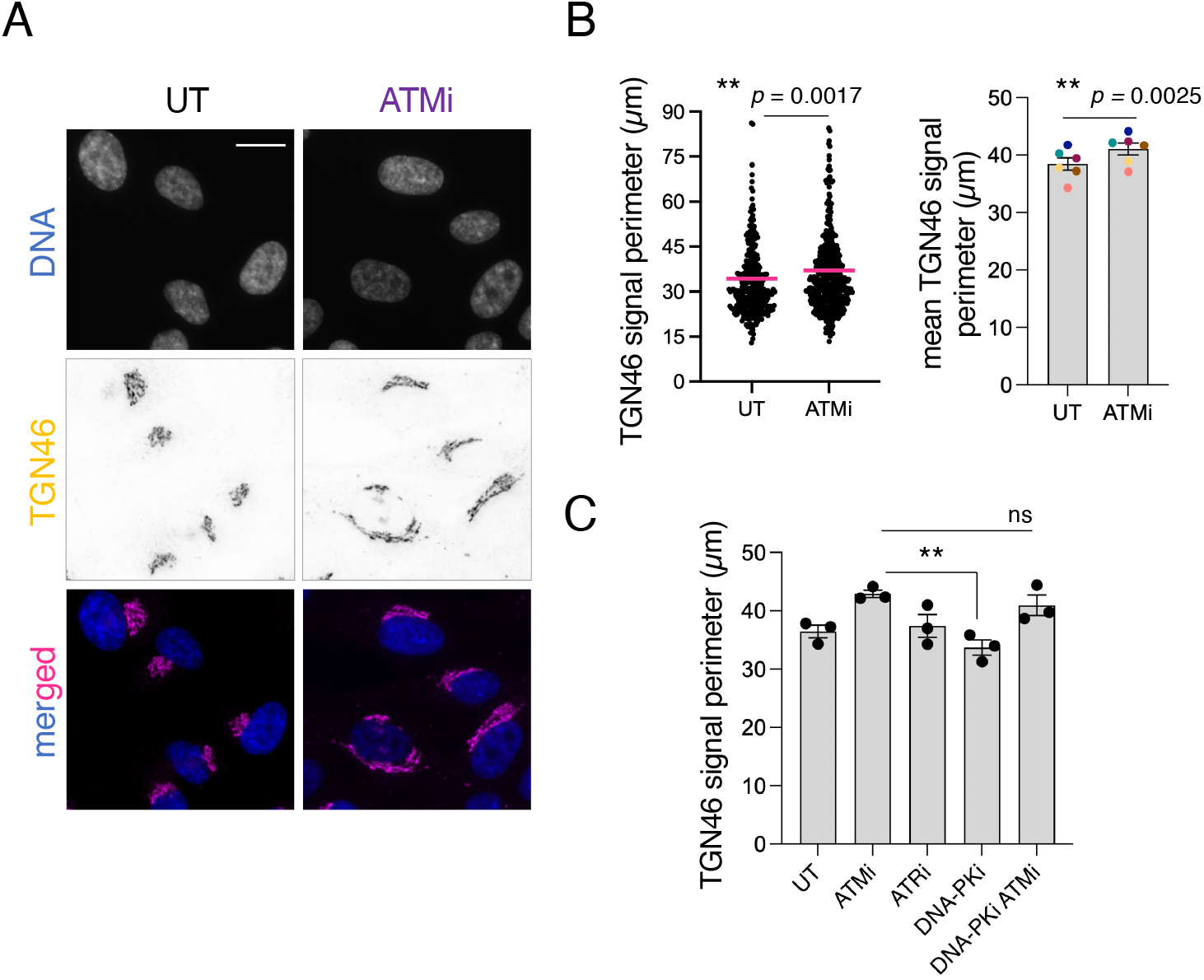
ATM kinase activity is important for Golgi shape. **A**.Immunofluorescence of the *trans* Golgi marker TGN46 in cells either left untreated (UT), or exposed to ATM inhibition using 500 nM AZD0156 for 4h. In the merged images, DNA is displayed in blue. Size bar is 12 μm. **B**.Quantification for the experiment shown in (A). **Left:** the individual values for the perimeter of individual Golgi unit signals (each point corresponds to one cell) as measured using TGN46 signals from one experiment were plotted for the indicated conditions. The horizontal pink bar indicates the mean of the population. The statistics result from applying a non-parametric Mann-Whitney test comparing the medians of the indicated samples with respect to the untreated condition. ns, non-significant. **Right:** the means from six independent experiments (color dots) as the one presented on the left were used to assess the reproducibility of the phenotypes for the indicated conditions. The bar height indicates the mean of the means and the error bars reflect the SEM. The statistics reveal the outcome of a paired *t*-test comparing the mean value between the two indicated conditions. **C**.As in (B, right) but for the indicated conditions. There is no color code because the replicates were not, in all cases, paired with each other. The “ATMi condition” mean was compared to all others by applying an unpaired one-way ANOVA test.

### 3. ATM phosphorylates target proteins at the Golgi, which limits its deployment

We next explored the meaning of the contribution of ATM kinase activity to limit Golgi deployment. On the one hand, the kinase DNA-PK can implement Golgi extension by phosphorylating GOLPH3 in response to long-lasting DNA damage treatment (at least 17 hours, maximal effects detected after 4 days) (11). Thus, since ATM silencing was imposed for 48 h in our set-up, our results could indirectly stem from DNA-PK responding to a presumed basal DNA damage accumulating without ATM. Alternatively, ATM may be exerting its kinase activity directly on Golgi substrates to restrict its extension. In the first scenario, the extension of the Golgi seen upon ATM inhibition (which also prevents its activity in the nucleus) should disappear when inhibiting DNA-PK, which we did not observe (Figure 2C) despite controlling that both inhibitors worked fine (Figure S3A). In the second scenario, monitoring ATM kinase activity on Golgi substrates using an antibody against anti-SQ/TQ-P (ATM’s phosphorylation motif) would be expected to reveal positive signals. In agreement with this second possibility, we could observe SQ/TQ-P signals at the Golgi of a subpopulation of cells. Analysis of both SQ/TQ-P-negative and -positive Golgi subpopulations (Figure S3B, see categorization criteria) revealed phosphorylation activity preferentially onto deployed Golgi units (Figure 3B). At this stage, a plausible interpretation was that the Golgi extension seen basally upon loss of ATM kinase activity may stem from its inability to phosphorylate Golgi proteins.

**Figure 3.**
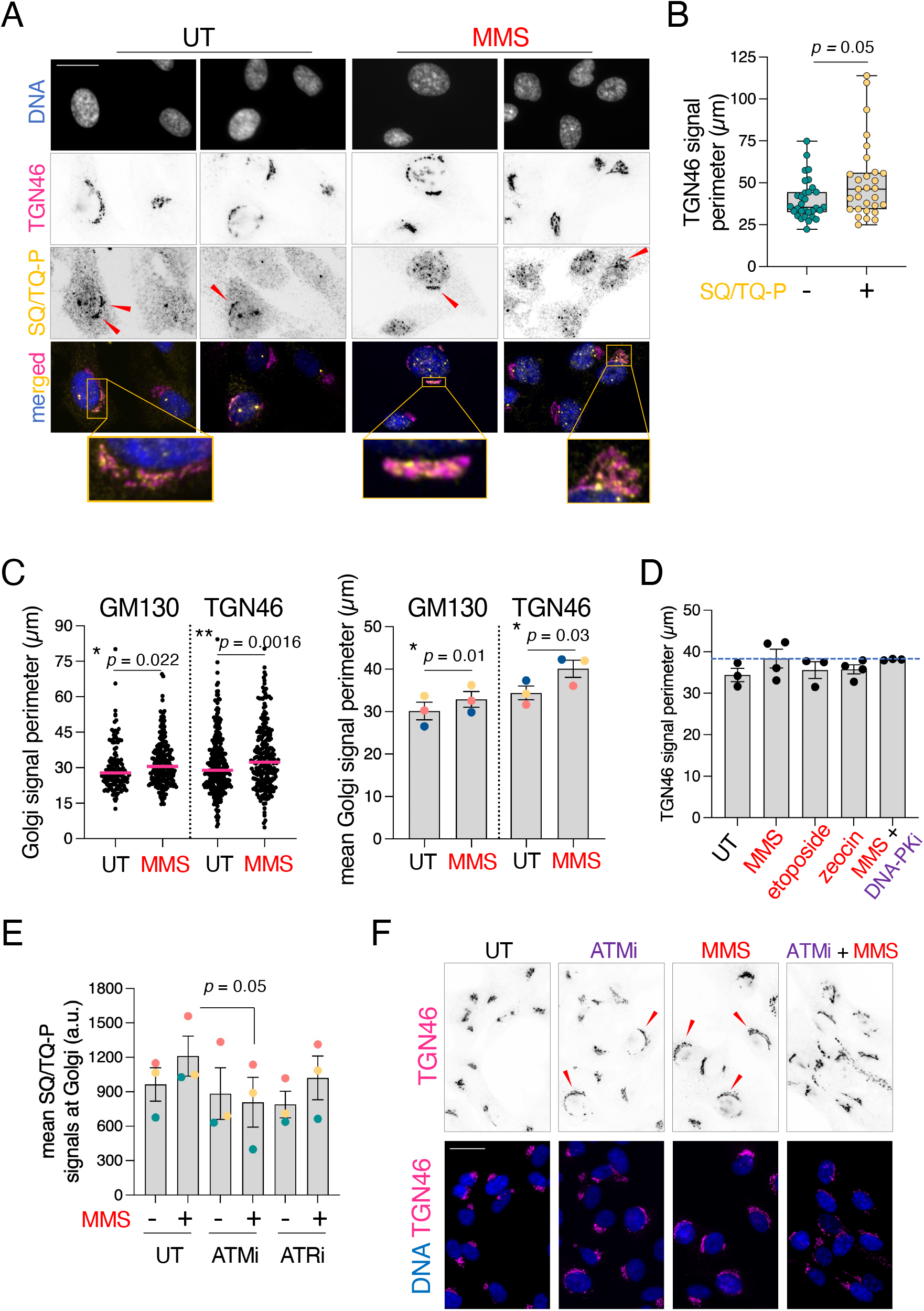
ATM kinase activity is detected onto Golgi units. **A**.Cells either untreated (UT) or exposed to MMS were processed for immunofluorescence to detect TGN46 and the target proteins phosphorylated by ATM and / or ATR on their SQ/TQ motif(s) (SQ/TQ-P). Two different examples are shown for each condition. Arrowheads point at SQ/TQ-P signals on the Golgi. Three regions have been zoomed out to better appreciate the spatial relationship of both signals (bottom). Size bar is 20 μm. **B**.Golgi units were ascribed to two categories, SQ/TQ-P-positive or -negative (criteria described in Figure S2B), and the perimeter of the units in both categories measured and plotted. n > 30 per category. A non-parametric Mann-Whitney test was applied to compare the medians of both distributions. **C**.Quantification experiments as shown in (A) with respect to TGN46 or GM130 signal perimeters in response to a 0.005% MMS treatment for 2h. **Left:** individual values for the perimeter of individual Golgi unit signals (each point corresponds to one cell) were plotted for the indicated conditions. The horizontal pink bar indicates the medians of the population. The statistics result from applying a non-parametric Mann-Whitney test comparing the medians of the treated sample with respect to the untreated condition. **Right:** the means from three independent experiments (color dots) as the one presented on the left were used to assess the reproducibility of the phenotypes for the indicated conditions. The bar height indicates the mean of the means and the error bars reflect the SEM. The statistics reveal the outcome of a paired *t*-test comparing the mean value between the two indicated conditions. **D**.Details as in (C) but for experiments that used the indicated molecules. There is no color code because the replicates were not, in all cases, paired with each other. The blue dashed line marks the level in the +MMS condition. **E**.Quantification of SQ/TQ-P signals detected at the Golgi of individual cells exposed to the indicated treatments. The means from three independent experiments (color dots) were used to assess the reproducibility of the phenotypes for the indicated conditions. The bar height indicates the mean of the means and the error bars reflect the SEM. The *p*-value stems from applying a paired t-test. **F**.Immunofluorescence of the *trans* Golgi marker TGN46 in cells either left untreated (UT), or exposed to ATM inhibition using 500 nM AZD0156 for 4h, or 0.005% MMS for 2h, or adding 500 nM AZD0156 first for 2h then, on top of that, 0.005% MMS for 2h. In the merged images, DNA is displayed in blue. Size bar is 20 μm.

To challenge this possibility, we aimed at provoking a strong Golgi deployment by creating DNA damage (11) and asking whether this triggered ATM kinase activity on the Golgi. We exposed cells to the genotoxin methyl methane sulfonate (MMS) and we confirmed it correctly induced the DDR by monitoring both nuclear ATM-P signals (Figure S3C) and nuclear SQ/TQ-P signals (Figure 3A). However, and in contrast with the report stating that DNA damage leads to a marked Golgi deployment assessed after (at least) 17 h of treatment (11), this short MMS treatment we applied to launch the DDR (i.e. 2h, which avoids altering cell cycle profiles) only led to a modest (though reproducible) increase in Golgi extension (Figure 3C). Treatment with other genotoxins (etoposide, zeocin) did not even elicit Golgi extension at all (Figure 3D). Furthermore, the MMS-induced Golgi extension was not abolished when inhibiting DNA-PK (Figure 3D), which would be expected for DNA damage-induced Golgi extension (11). These experiments reveal that DNA-PK phosphorylation-mediated Golgi extension is not an early event in the response to DNA damage. Of relevance, though, the Golgi extension provoked by MMS treatment matched a marked increase in SQ/TQ-P signals at the Golgi apparatus (Figure 3A, MMS, red arrowheads; Figure 3E), and was a rare but detectable event in response to other genotoxins, such as etoposide (Figure S3D). Since SQ/TQ-P signals can be due to either ATM or ATR activity (12), we asked on which of these kinases SQ/TQ-P signals depended, for which we pre-incubated the cells for 2h with specific inhibitors targeting each of them. The MMS-induced nuclear SQ/TQ-P signals were fully abolished without ATM activity, while they were unaffected without ATR activity (Figure S3E). The same was true for MMS-induced SQ/TQ-P signals at the Golgi (Figure 3E).

Last, the data made us postulate that, if ATM prevents excessive Golgi extension, combining ATM inhibition with MMS, both provoking its extension, should lead to an excessive deployment. In agreement, lack of ATM kinase activity during MMS treatment led to aberrantly fragmented Golgi signals (Figure 3F, ATMi + MMS). It is worth acknowledging that ATM inhibition during MMS treatment led to a detectable decrease in ATM total levels (Figure S2A, MMS), which may add up to the strength of the phenotype. Altogether, we conclude that ATM restricts an excessive deployment of Golgi units by exerting its kinase activity on Golgi targets. This occurs both basally and upon cues that further stimulate the ability of ATM to probe membranes’ status (see discussion), for example MMS.

### 4. ATM control of Golgi stretching can impact Golgi client protein maturation

Golgi function relies on Golgi morphology and, as such, GOLPH3/MYO18A/F-actin-dependent Golgi deployment is critical for Golgi protein cargo maturation and subsequent trafficking to the plasma membrane (10, 13, 14). In agreement, in the absence of GOLPH3, cargo protein glycosylation patterns are perturbed (15). TGN46, heavily modified by glycosylation itself (16), acts as a crucial sorter for a specific class of vesicles mediating *trans* Golgi to plasma membrane traffic (17). Despite displaying a steady-state localization at the *trans* Golgi (this is why it is commonly used as a *trans* Golgi marker), TGN46 rapidly cycles between the *trans* Golgi and the plasma membrane (18), accompanying the cargoes it helps to sort, therefore virtually behaving as a cargo itself.

In view of our findings, we explored if ATM absence could impact Golgi performance during cargo protein modification, for which we exploited TGN46 as a readout. We observed no significant modification of its glycosylation pattern in the absence of ATM (Figure 4A). On the contrary, GOLPH3 absence, which prevents Golgi spreading, led to an alteration of TGN46 electrophoretic pattern, as anticipated, with an increase in the presence of the hyper-modified form (Figure 4A, “H”). We asked if the increased Golgi deployment promoted by ATM absence in this genetic background could restore, at least partly, normal-looking TGN46 glycosylation profiles. When we combined the double depletion of GOLPH3 with ATM, the TGN46 glycosylation profile resembled that of the controls (Figure 4A). Thus, the Golgi deployment imposed by ATM absence can affect in-Golgi protein maturation.

**Figure 4.**
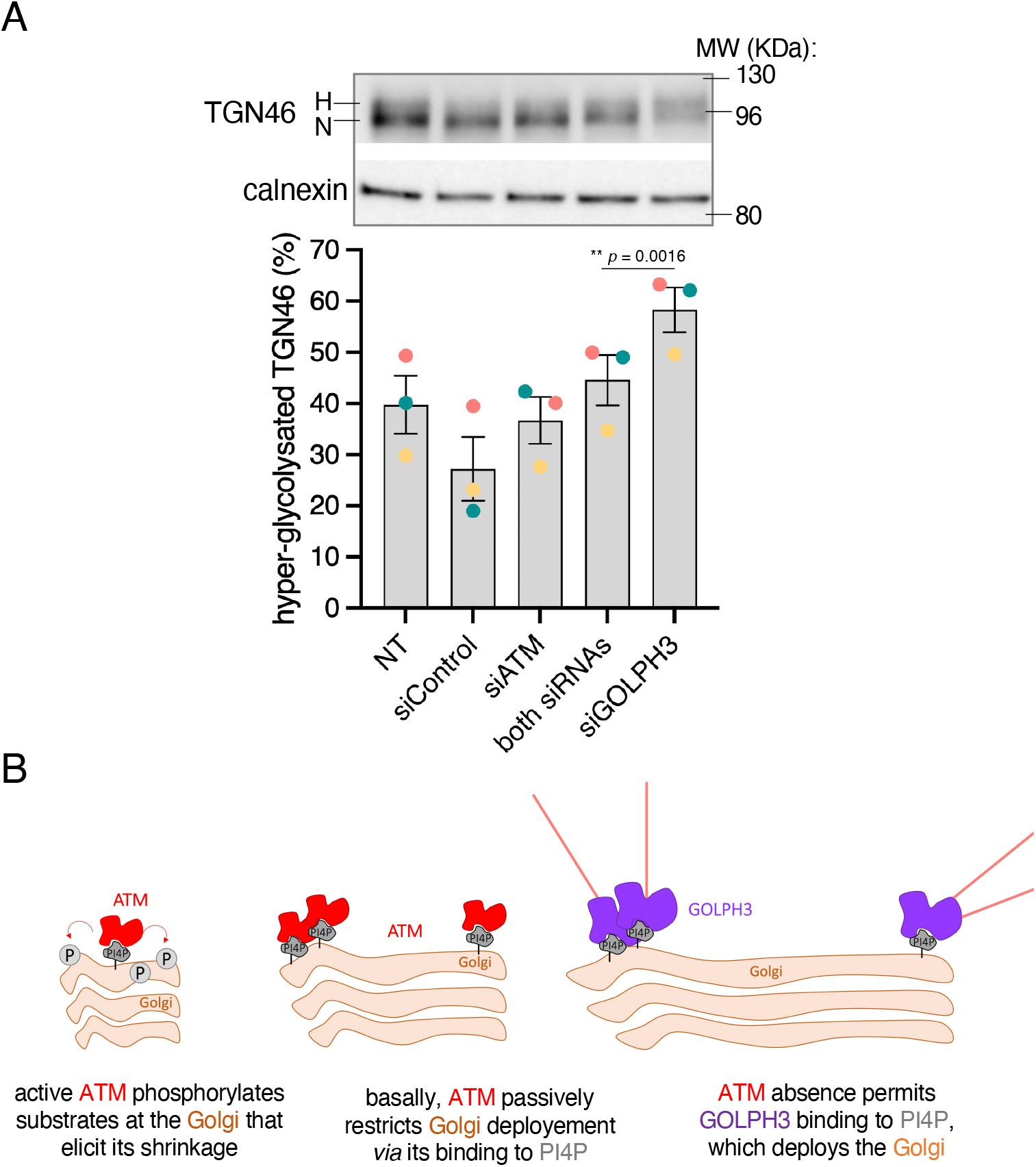
ATM absence impacts Golgi activities. **A.Top:** Western blot to monitor the status of TGN46 glycosylation (H = hypermodified, upper band; N = normal glycosylation, bottom band) in the indicated conditions. Calnexin is shown to monitor the total protein load. **Bottom:** the percentage of hypermodified TGN46 was calculated from western blots from three independent experiments and plotted as three different color dots. The bar height indicates the mean of the means and the error bars reflect the SEM. The statistics reveal the outcome of a paired *t*-test comparing the mean between the two indicated conditions. **B**.Model: ATM passively contributes to Golgi morphology thanks to its basal binding to Golgi-resident PI4P (middle). In its absence, exposed PI4P can be bound by GOLPH3, which stretches the Golgi, leading to an extended morphology (right). Additionally, ATM can use its kinase activity to phosphorylate Golgi-resident or Golgi-proximal substrates, which actively contribute to maintain a compact Golgi morphology (left).

In sum, we propose that ATM residence at the Golgi apparatus is not anodyne, for it serves to check up on Golgi shape, likely by virtue of its binding to PI4P and / or by competing with GOLPH3 for this binding, either passively, and / or through an inactivating phosphorylation. In doing so, ATM restricts excessive Golgi deployment, both basally and in response to challenges, thus warranting proficient Golgi activity, such as cargo protein maturation.

## Discussion

In this work, we report that the kinase ATM, well known for its role in signaling DNA damage in the nucleus, is important to ensure the homeostasis of the Golgi. Its absence entails a deployment of this organelle, which in part can be attributed to GOLPH3. GOLPH3 and ATM, both being PI4P binders, and GOLPH3 activity in Golgi stretching being mediated by its binding to PI4P, we presume that ATM absence permits GOLPH3 to better access this lipid. On top of that, we detect that ATM can exert its kinase activity onto Golgi substrates, and that failure to do so also leads to excessive Golgi deployment in various cell lines of human origin, suggestive of a general phenomenon. The “kinase mode” of control occurs under basal conditions and becomes more prevalent whenever ATM is stimulated by cues such as MMS treatment.

Importantly, the morphological alterations displayed by the Golgi whenever ATM is not functional can alter Golgi cargo biology.

### Antagonistic roles for ATM and DNA-PK on Golgi morphology

When we used the genotoxin MMS, we monitored a consistent deployment of the Golgi, although in response to short treatment and a genotoxin not tested before. Previous works assessed the impact of camptothecin, doxorubicin and of ionizing radiation (11, 19). At that time, it was found that the DNA damage-responding kinase DNA-PK mediated Golgi deployment in response to these challenges by phosphorylating GOLPH3 on its T143, which would enhance its interaction with the stretching tandem MYO18A/F-actin. The potential role of ATM on mediating Golgi deployment was tested and the authors concluded that absence of ATM did not mediate such an event (11). In strong agreement with those findings, we now confirm that ATM role is not to promote Golgi extension, for it is to prevent it. We find that ATM absence leads to a more deployed Golgi, that ATM exerts its phosphorylation onto Golgi entities with deployed aspect, and that restriction of ATM kinase activity matches an increase in Golgi size. The DNA-PK complex, composed of the DNA-binding Ku70/80 heterodimer and the catalytic subunit DNA-PKcs, is the key regulator of DSB repair through non-homologous end joining. ATM and DNA-PK both become activated by, and contribute to handle (though differently) harms such as DNA breaks, oxidative stress and hypoxia (reviewed in (20)). Our discovery implies that ATM and DNA-PK exert antagonistic roles, and this with different kinetics, likely orchestrating a two-component response: in response to a challenge, ATM may ensure Golgi shape conservation early after the stress, while later extension is ensured by DNA-PK. This places these two DDR master kinases on the two sides of a balance ensuring Golgi homeostasis.

### ATM impacts Golgi function

Golgi morphology is not a necessarily accurate predictor of its performance (21), since the impact exerted by a manipulation greatly depends on the specific mechanism of the perturbation. However, it is established that, for perturbations of the GOLPH3 pathway, the consequences to Golgi morphology and secretory function are predictable. Indeed, while a moderate increase in GOLPH3 pathway activity leads to an increase in Golgi-mediated anterograde trafficking, Golgi fragmentation out of a hyper-active GOLPH3 pathway leads to impaired trafficking (11, 14). Our work suggests that, at least in part, the effect of ATM absence on Golgi morphology is mediated by GOLPH3 (Figure 1). Mild overexpression of GOLPH3 leads to an extended Golgi ribbon, while very strong overexpression results in such an extension that it ends up in fragmentation (14). This is reminiscent to the different stages we observe, where ATM absence alone, or MMS treatment, only modestly deploy the Golgi, but the combination of both interventions ends up in Golgi fragmentation (Figure 3F). It appears therefore predictable that alterations in cargo glycosylation arising as a consequence of GOLPH3 absence may be palliated if Golgi morphology is restored. In this sense, we monitored the normalization of glycosylated TGN46 when GOLPH3 absence is compensated by lack of ATM (Figure 4A). Thus, the crosstalk between ATM and GOLPH3 at the Golgi is likely to condition functional consequences for Golgi-driven transport. Of note, while our visual inspection of Golgi signals revealed that, upon ATM inhibition, some units were markedly deployed (Figure 2A), the quantification showed only a small mean increase in Golgi perimeter (3 μm). This implies a non-negligible heterogeneity within the population, and suggests that not all Golgi entities are equally susceptible of being controlled by ATM. Plausible variables conditioning this proneness and that deserve further exploration could be the cell cycle stage, or the metabolic status.

### Activation modes for Golgi-resident ATM and its target substrates

The Golgi is, after the microtubule-organizing center (MTOC), the second major node for microtubule nucleation (22), given that membrane components of the Golgi can also nucleate and stabilize microtubules at both the *cis* and *trans* Golgi (23, 24). In response to DNA-damaging agents, a drastic microtubule reorganization that leads to their stabilization at the Golgi takes place (25). ATM is reported to exert its kinase activity in the cytoplasm onto cytoskeletal targets in response to mechanical stress (26). It is therefore very tempting to propose, in view of our data, that the stabilization of microtubules at the Golgi and the ATM-dependent phosphorylation of cytoskeletal targets, both occurring in response to stress, concern the same phenomenon, orchestrated by the Golgi-based ATM kinase activity that we report here. But which would be the targets of this ATM kinase activity? Our data open the possibility that GOLPH3 is one of them. Among ATM’s cytoskeletal targets, there were myosin1B, TUBB3, leiomodin-1, zyxin, granulin (a Golgi cargo), and DENND2A (likely involved in retrograde transport from endosomes to Golgi) specifically in response to mechanical stress (26). It is highly possible that ATM substrates may vary depending on the activating stress.

In this sense, ATM activation, as defined by its autophosphorylation, has been reported in response to multiple cues, many of which are not necessarily linked to DNA integrity, or even the nucleus, such as chromosome condensation, hyperthermia and hypoxia (27–32). In response to DNA double strand breaks (DSBs), the MRE11-RAD50-NBS1 (MRN) complex recruits ATM to the lesion and activates it (33, 34). Under oxidative stress conditions, such as H_2_O_2_ treatment, ATM activation does not depend on the MRN complex but on ATM homo-dimerization through intermolecular disulfide bonds involving the critical cysteine residue Cys2991 (35). Of note, ATM substrate phosphorylation is different between oxidative stress-induced ATM-mediated phosphorylation and MRN/DSB-dependent ATM-imparted phosphorylation (35). ATM substrates in response to oxidative stress do not appear to be enriched in Golgi-related hits, but mostly on proteostasis and mitochondrial function (reviewed in (36)). Of relevance, upon genotoxic attack, a battery of DNA repair-related proteins, discovered to basally reside on the Golgi apparatus, relocate to the nucleus, where they support repair, in an ATM-dependent manner (37). This suggests that Golgi-resident ATM may phosphorylate them to instruct their mobilization. It remains unclear, in this latter case, how Golgi-based ATM becomes active then modifies substrates at the Golgi in response to a signal that emanates from elsewhere in the cell, namely damaged DNA within the nucleus. One option would be that DNA-PK, activated at nuclear DNA breaks, phosphorylates GOLPH3, engaging it in Golgi deployment, as reported (11). This physical stretching of the Golgi would constitute the signal activating Golgi-resident ATM. However, we showed here that genotoxins very poorly induce Golgi extension during short treatments. Alternatively (and not exclusively), physical cues at the Golgi could stem from altered nuclear mechanics, consequence of chromatin alterations (38), that would propagate to adjacent membranous systems (39). It still remains unanswered how ATM would “feel” membrane tension, although its numerous HEAT repeats (40) were already postulated to likely act as a membrane sensor (26). In further support, we already reported that MMS, beyond its genotoxic action, specifically elicits membrane stress (41), which may help explain why we find it triggers a Golgi deployment that other genotoxins do not (Figure 3).

In sum, we propose that ATM passively contributes to Golgi morphology thanks to its basal binding to Golgi-resident PI4P (Figure 4B, middle). In its absence, exposed PI4P can be bound by GOLPH3, which stretches the Golgi, leading to an extended morphology (Figure 4B, right). Additionally, ATM can use its kinase activity to phosphorylate Golgi-resident or Golgi-proximal substrates, which actively contribute to maintaining a compact Golgi morphology (Figure 4B, left). In this sense, ATM inactivation in combination with any cue instructing Golgi extension may end up in Golgi fragmentation. We believe that this work broadens our understanding of Golgi biology and enforces the notion that different organelles, such as the Golgi and the nucleus, coordinate with each other.

## Material and Methods

### siRNAs and antibodies

siControl was the universal negative control siRNA #1 (SIC001, Sigma-Aldrich) siATM (SASI_Hs01_00093615, 5’ -CUUAGCAGGAGGUGUAAAU [dT][dT]-3’, Sigma-Aldrich) siGOLPH3 (SASI_Hs01_00133693, 5’-GGUGUAUUGACAACAGAGA[dT][dT]-3’, Sigma-Aldrich) **Primary antibodies**: Anti-PI4P (Z-P004, Echelon Biosciences, IF at 1/100); anti-GOLPH3 (SAB4200341, Sigma-Aldrich, IF at 1/300, WB at 1/2,000); anti-TGN46 (AHP500G, Bio-Rad, IF at 1/300, WB at 1/2,000); anti-GM130 (HPA021799, Sigma-Aldrich, IF at 1/300); anti-P-Ser1981-ATM (4526, Cell Signaling, IF at 1/200); anti-ATM (GTX132147, CliniSciences, WB at 1/2,000); anti-SQ/TQ-P (2851S, Ozyme, IF at 1/500); anti-PGK1 (13472727, ThermoFisher, WB at 1/1,000); anti-Calnexin (610523, Becton Dickinson, WB at 1/1,000); anti-_Y_H2AX (05-636, Millipore, IF at 1/500). **Secondary antibodies** for WB: anti-rabbit (10733944, Fisher Scientific, WB at 1/15,000); anti-mouse (A9044, Sigma-Aldrich, WB at 1/5,000); anti-sheep (A3415, Merck, WB at 1/2,500). For IF, all used at 1/500: anti-rabbit (406406, Biolegend), anti-mouse (A-21123, Fisher Scientific), anti-sheep (A-21099, Thermo Fisher).

### Cell culture, transfection and treatment

RPE-1 and Huh-7 cell lines used in this study were authenticated by ATCC STR profiling. A549 cells were a kind gift from Caroline Goujon, originally obtained from the ATCC (ATCC CRM-CCL-185). Cells were grown in DMEM Glutamax (31966047, Gibco) supplemented with 10 % FBS (F7524, Merck) and 1 % Pen/Strep (15140122, Gibco) at 37°C and 5% CO_2_, they were split using trypsin solution (0.25% trypsin (15090046, Gibco), 0.53 mM EDTA pH8, in 1X PBS) and were seeded to be subconfluent the day of harvesting. Medium was changed before applying the intended treatments, cells were collected for western blot or fixed on coverslips for immunofluorescence. For transfections, cells were seeded 24 h prior to transfection with Lipofectamine RNAiMAX (13778075, Invitrogen). The supplier’s protocol was followed for 10 nM siRNA and 48 h of transfection. Pretreatments with the inhibitors were applied for a total of 4h, AZD0156 (ATMi, HY-100016, Clinisciences) at 0.5 μM; VE-821 (ATRi, HY-14731, Clinisciences) at 5 μM; NU7026 (DNA-PKi, A12752-10, Clinisciences) at 5μM. Genotoxins were always applied for 2h, at the following doses: MMS (129925, Sigma-Aldrich) at 0.005 %; etoposide at 1μM; zeocin (10072492, Fisher scientific) at 10 μg/mL.

### Immunofluorescence

Cells fixed 20 min with 4% PFA/1X PBS, except if for P-ATM detection (10 min with cold 100% methanol in that case), then coverslips were washed once with 1X PBS. For permeabilization, 0.2% Triton/1X PBS for 10 min was applied, except if subsequent PI4P detection (100 μM digitonin/1X PBS for 5 min in that case), or downstream P-ATM detection (step skipped in this case). Blocking was done with 3% BSA/1X PBS for 30 min. Coverslips were incubated with primary antibodies diluted in 3% BSA/1X PBS for 1 h and then washed 3 times 1X PBS under gentle shaking, then incubated with secondary antibodies diluted in 3% BSA/1X PBS supplemented with 1 μg/mL DAPI (D9542, Sigma-Aldrich) for 1 h, then washed 3 times with 1X PBS under gentle shaking and washed three times with H_2_O. Finally, coverslips were allowed to dry and mounted using ProLong Gold Antifade (P36930, ThermoFisher) and then left to harden overnight at room temperature in the darkness. Acquisitions were done using epifluorescence microscopy with an upright microscope (Zeiss AxioImager Z2), at x40 magnification. The precise details for how Golgi signal perimeter and circularity were selected and analyzed are explained in Annex 1. Quantification of nuclear signals was done delimiting nuclei as region of interest (ROI) *via* thresholding, applying this ROI onto the other channel of interest (_Y_H2AX or SQ/TQ-P) and measuring the mean intensity. To quantify SQ/TQ-P signals at the Golgi, a similar macro was used but that delimits first the Golgi area (using a Golgi marker) *via* an intensity threshold, then applying it onto SQ/TQ-P channel, in the end measuring the mean SQ/TQ-P intensity per “Golgi unit”.

### Western Blot

Samples were lysed with high-salt buffer: 50 mM Tris pH 7.5, 300 mM NaCl, 1% Triton x100, mini cOmplete protease inhibitor cocktail (11836153001, Roche), Halt Phosphatase inhibitor cocktail (78420 ThermoFisher), 100 μL per well of a 6 well plate, incubated on ice for 10 min with frequent vortexing. Samples were centrifuged for 10 min at 17,000 *g* at 4°C, and supernatants quantified using the Pierce™ BCA kit (10741395, ThermoFisher). For subsequent western blot, 20-30 μg of whole-cell extracts were loaded onto precast 4-20% acrylamide gels (5671094, BioRad) and migrated 70 min at 150 V in migration buffer (25 mM Tris, 200 mM Glycine, 0.1% SDS). The proteins were transferred to a nitrocellulose membrane using Transblot Turbo system (BioRad) with “mixed weight” program, then blocked with 5% milk/TBS-T for 30 min. Incubation with primary antibodies was done in antibody solution (3 % BSA, 0.01 % Sodium Azide in 1x TBS-Tween_0.1%,_) for 1 h, washed 3 times 5 min in 1x TBS-Tween_0.1%,_ incubation with secondary antibodies in 3% BSA/1x TBS-Tween_0.1%,_ for fluo-coupled antibodies, or 5% milk/1x TBS-Tween_0.1%,_ for HRP-coupled antibodies for 1h, washed 3 times 5 min in 1x TBS-Tween_0.1%,_. For fluo-coupled antibodies, membranes were dryed between two Whatmann’s paper; for HRP-coupled antibodies SuperSignal™ West Pico PLUS was used, and imaging done with Odyssey® M Imager (LI-COR). To quantify the signal emanating from hypermodified TGN46, using FIJI, a box was drawn per lane that would contain all TGN46 signals, the area of such signals quantified and ascribed either to normal (N, bottom part of the box) or hypermodified (H, upper part of the box). The percentage of “H” signal with respect to the total was used for the plots.

### Cytometry

3 × 10^6^ cells were retrieved after trypsinization then pelleted by centrifugation 5 min at 200 *g*, resuspended in 1 mL PBS 1X, then 2.5 mL of 100 % ethanol were added and vortexed well. Incubation at -20°C for 30 min was followed by resuspension with 5 mL of PBS 1X, centrifugation for 5 min at 500 *g*, and resuspension of the pellet in 1 mL 50 μg/mL RNAseA/1X PBS and incubation at RT for 30 min. Cells were then subjected to centrifugation at 200 *g* for 5 min, resuspended in 1 mL 2N HCl solution, incubated at RT for 30 min, then 3 mL of 1X PBS were added. Centrifugation 5 min at 200 g was followed by pellet resuspension in 1 mL 0.1M borate solution, incubation at RT for 2 min 30 sec, then addition of 3 mL 1X PBS. Centrifugation at 200 *g* 5 min, resuspension of the pellet in 200 μL of 1X PBS and addition of propidium iodide at 25 μg/mL final, incubation at least 10 min at RT. Acquisition and analysis were done using Novocyte Express.

### Data plotting and analysis

GraphPad Prism was used to plot all the graphs and to statistically analyze the data. For statistical analyses, the choice of test was justified by the nature of the sample and is explained for each particular case within the figure legends. Whenever nothing is indicated on the graphs, differences are not significant (ns).

## Abbreviations

ATM: ataxia telangiectasia-mutated kinase
ATR: ataxia telangiectasia and Rad3-related kinase
DDR: DNA damage response
DNA-PK: DNA-dependent protein kinase
MMS: methyl methane sulfonate
MTOC: microtubule organizing center
PIKK: phosphatidyl- inositol-3-kinase-related kinase
PI4P: phosphatidyl-inositol-4-phosphate
SEM: standard error of the mean

## Authors’ contributions

Conceptualization: M.M.-C.; Data curation: C. S.; J.C.; M.M.-C.; Formal analysis: C. S.; J.C.; M.M.-C.; Funding acquisition: M.M.-C.; Investigation: C. S.; J.C.; Methodology: C. S.; M.M.-C.; Project administration: M.M.-C.; Supervision: C. S.; M.M.-C.; Visualization: C. S.; J.C.; M.M.-C.; Writing – original draft: C. S.; M.M.-C.; Writing – review & editing: J.C.; C. S.; M.M.-C.

## Acknowledgments

We acknowledge the imaging facility MRI, member of the national infrastructure France-BioImaging, supported by the French National Research Agency (ANR-24-INBS-0005 FBI BIOGEN). We thank the French National Research Agency (ANR) for supporting this work *via* two different grants: “Genome Sterolity” (ANR-21-CE12-0004-01) and “PATHaFIX” (ANR-25-CE15-5302).

## Supplementary Figures

**Figure S1.**
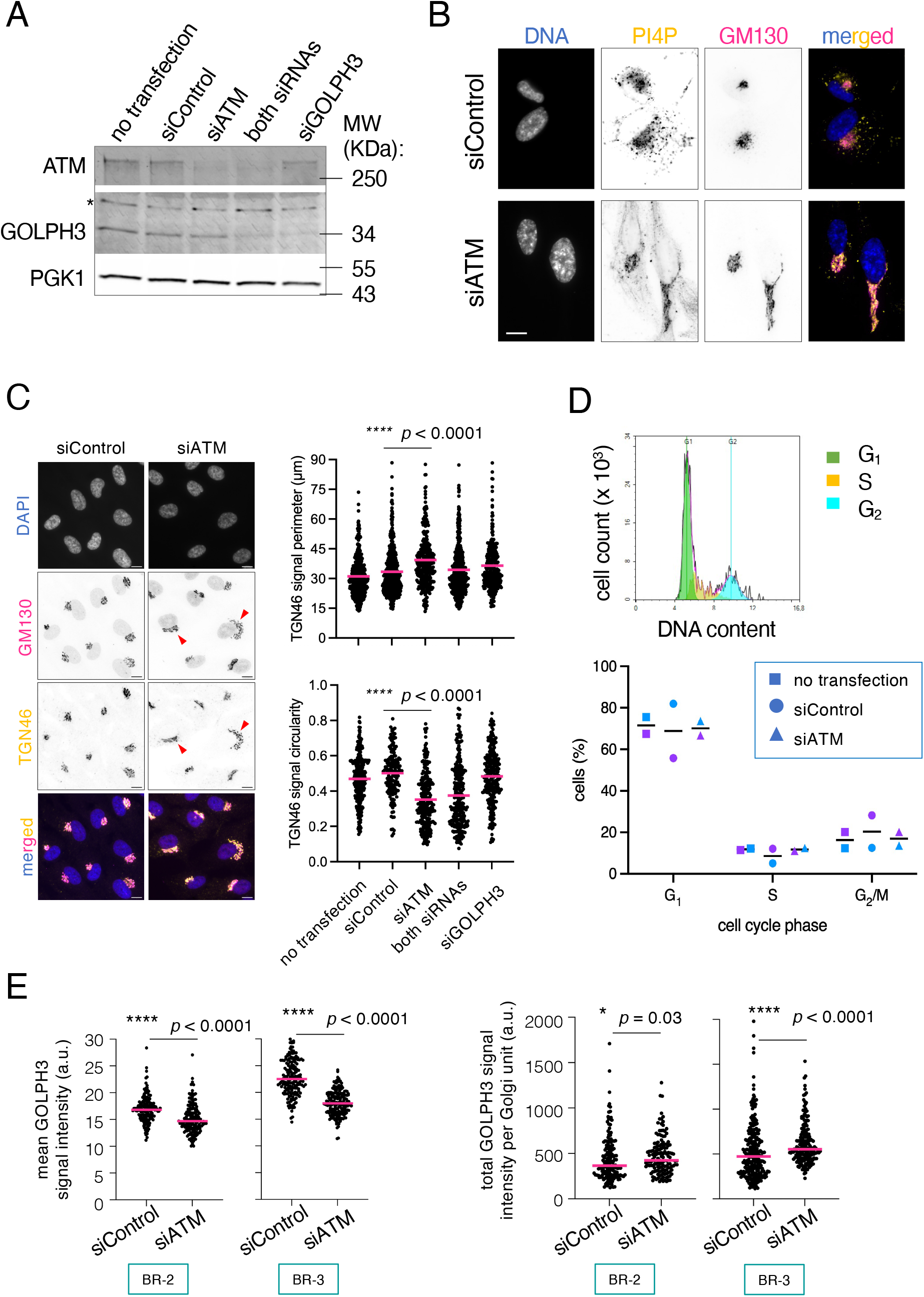
Support information for Figure 1. **A**. Western blot of the indicated proteins to monitor the efficiency of the siRNA-mediated depletion of ATM and GOLPH3. The asterisk indicates an unspecific band. PGK1 is included as a loading control. **B**. Immunofluorescence of the *cis*-Golgi marker GM130 and the lipid PI4P in cells transfected either with siControl or siATM. Size bar is 10 μm. **C**. Immunofluorescence of the *cis*-Golgi marker GM130 and the *trans* Golgi marker TGN46 in cells transfected either with siControl or siATM, as in Figure 1A, accompanied by the quantification displaying the individual values for the perimeter (top) and the circularity (bottom) of individual Golgi unit signals (each point corresponds to one cell) as measured using TGN46 signals from one experiment. The horizontal pink bar indicates the mean of the population. Arrowheads in the pictures point to deployed Golgi units. The statistics result from applying a non-parametric Mann-Whitney test comparing the medians of the indicated samples. Size bar is 5 μm. **D. Top:** prototypical profile of cytometry assessments, in which the number of cells (y-axis) is plotted against their DNA content (x-axis), and ascribed to a specific cell cycle phase (colors) by the software. **Bottom:** The percentages of cells present in each cell cycle phase were measured and plotted for the indicated conditions out of two independent transfection experiments (blue, violet). The black horizontal bar indicates the mean of those experiments. **E**. Two additional, independent biological replicates (BR) of the experiment presented in Figure 1D. **Left:** quantification of the mean GOLPH3 signal intensity per pixel and **Right:** of total GOLPH3 signal intensity in each complete Golgi unit out of one individual experiment. Each dot corresponds to a Golgi unit from an individual cell. The horizontal pink bar indicates the median of the population. The statistics result from applying a non-parametric Mann-Whitney test comparing such medians.

**Figure S2.**
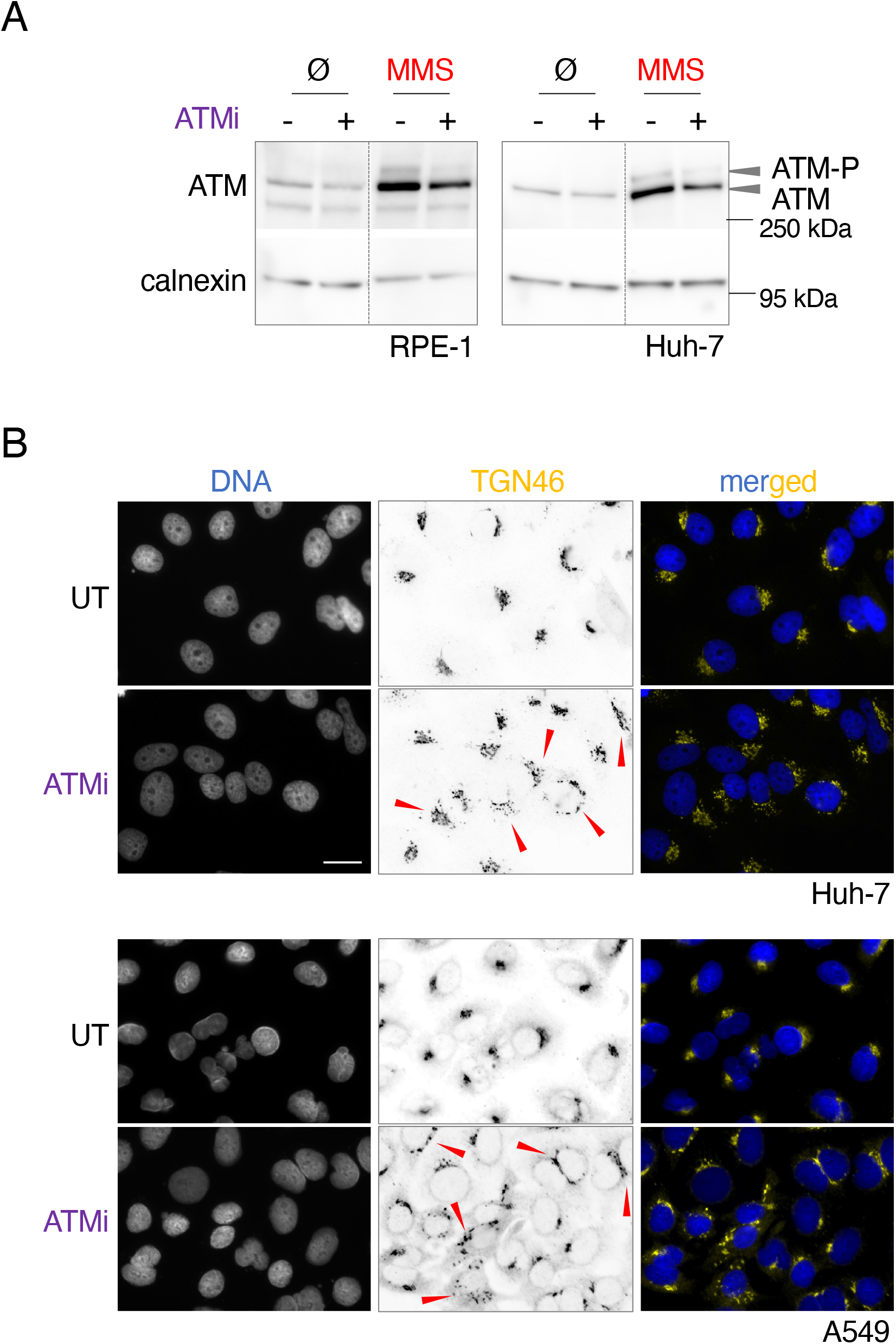
Support information for Figure 2. **A**. Western blot of total levels of ATM in the indicated cell lines from cells left untreated (UT), or exposed to ATM inhibition using 500 nM AZD0156 for 4h. The images shown for RPE-1 cells were already presented by us in (9). Arrowheads highlight the non-phosphorylated and the phosphorylated isoforms that can be detected. Calnexin is used as a loading control. **B**. Immunofluorescence of the *trans* Golgi marker TGN46 in the two indicated cell lines. Cells were either left untreated (UT), or exposed to ATM inhibition using 500 nM AZD0156 for 4h. In the merged images, DNA is displayed in blue. Size bar is 15 μm.

**Figure S3.**
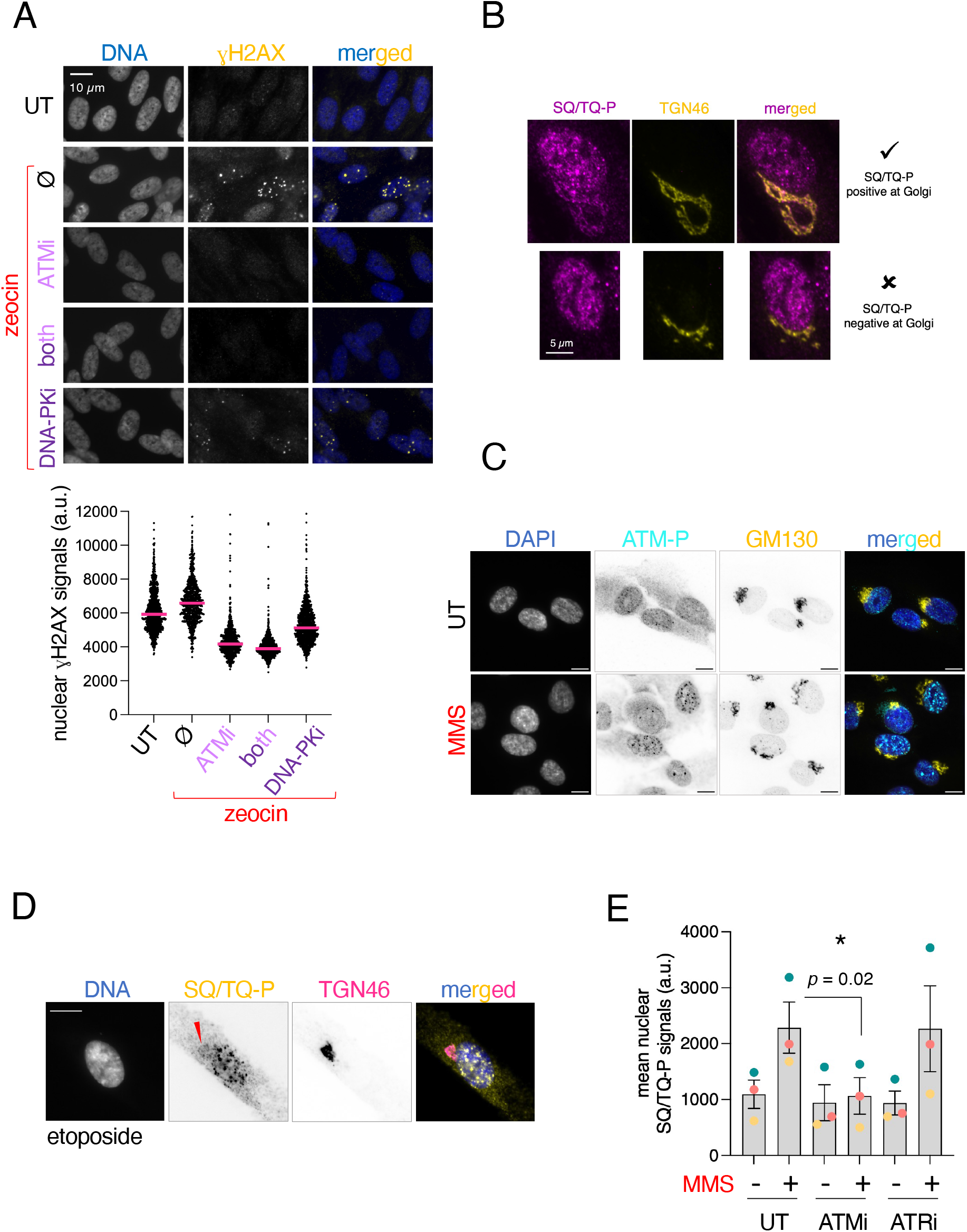
Support information for Figure 3. **A. Top:** Immunofluorescence of _Y_H2AX in response to DNA damage (zeocin) on RPE-1 cells that had been kept under confluence for 2 weeks, the idea being that the response to DNA damage is mostly mediated by ATM and DNA-PK in non-cycling cells (42). The goal of this experiment was to control the efficiency of ATM and of DNA-PK inhibitors. **Bottom:** values obtained for _Y_H2AX nuclear signals (each point corresponds to one nucleus) from one experiment were plotted for the indicated conditions. The horizontal pink bar indicates the median of the population. **B**. Representative images illustrating the criteria followed in Figure 3B to ascribe Golgi units to the two considered categories: either SQ/TQ-P-positive (top) or -negative (bottom). **C**. Immunofluorescence of the phosphorylated form of ATM (ATM-P-Ser1981) and the *cis*-Golgi marker GM130 in cells either left untreated (UT), or exposed to 0.005% MMS for 2h. The size bar is 5 μm. **D**. Immunofluorescence to detect SQ/TQ-P signals. The example frames one cell that displays co-localization with the Golgi marker TGN46. Size bar is 10 μm. **E**. Quantification of SQ/TQ-P signals detected within the nucleus of individual cells exposed to the indicated treatments. The means from three independent experiments (color dots) were used to assess the reproducibility of the phenotypes for the indicated conditions. The bar height indicates the mean of the means and the error bars reflect the SEM. The *p*-value stems from applying a paired t-test.

**Annex 1.**
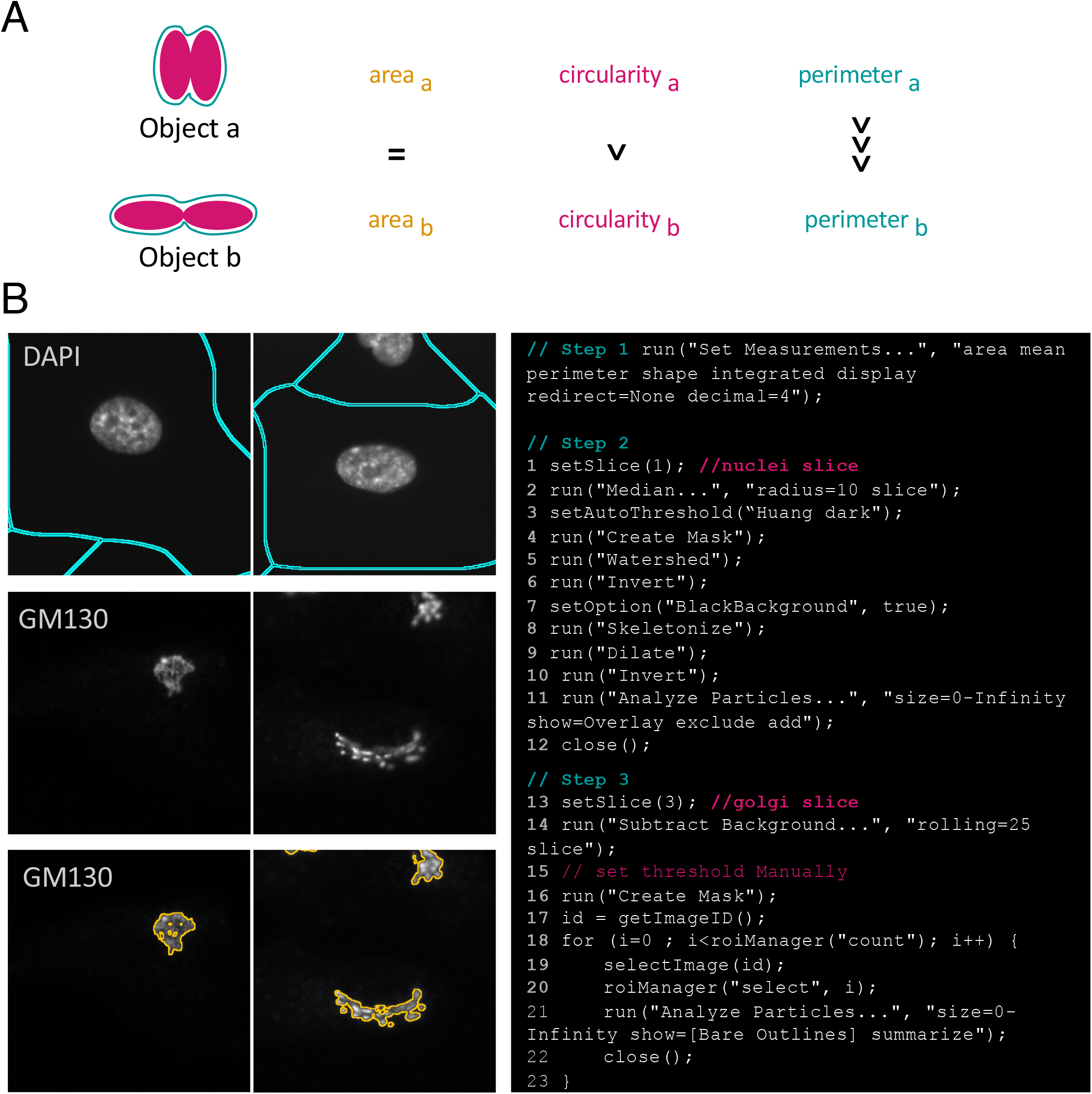
**A**.Rationale behind the choice of perimeter and circularity as appropriate readouts for quantifying Golgi morphology, especially important for objects of very similar area. **B**. Procedure for quantifying perimeter and circularity in a semi-automated manner using FIJI macro’s mode: Step 1 is the choice of adequate values to measure; Step 2 allows to create a fictive cytoplasm based on the position of nuclei; Step 3 establishes a threshold, as selected by the user through visual inspection, capable of faithfully capturing the whole individual Golgi unit under consideration (GM130 signals in this example). Left is visual illustration of the steps; right is the actual macro syntax.

## References

1. A. N. Blackford, S. P. Jackson, ATM, ATR, and DNA-PK: The Trinity at the Heart of the DNA Damage Response. Mol. Cell 66, 801–817 (2017).

2. T. Hunter, When is a lipid kinase not a lipid kinase? When it is a protein kinase. Cell 83, 1–4 (1995).

3. Z. Cui, A. Esposito, G. Napolitano, A. Ballabio, J. H. Hurley, Structural basis for mTORC1 activation on the lysosomal membrane. Nature 647, 536–543 (2025).

4. C. Yi, et al., Formation of a Snf1-Mec1-Atg1 Module on Mitochondria Governs Energy Deprivation-Induced Autophagy by Regulating Mitochondrial Respiration. Dev. Cell 41, 59-71.e4 (2017).

5. A. Kumar, et al., ATR mediates a checkpoint at the nuclear envelope in response to mechanical stress. Cell 158, 633–646 (2014).

6. X. H. Zhang, C. Zhao, Z. A. Ma, The increase of cell-membranous phosphatidylcholines containing polyunsaturated fatty acid residues induces phosphorylation of p53 through activation of ATR. J. Cell Sci. 120, 4134–4143 (2007).

7. J. Luessing, C. C. Okowa, E. Brennan, M. Voisin, N. F. Lowndes, A function for ataxia telangiectasia and Rad3-related (ATR) kinase in cytokinetic abscission. iScience 25, 104536 (2022).

8. S. Ovejero, C. Soulet, S. Kumanski, M. Moriel-Carretero, Coordination between phospholipid pools and DNA damage sensing. Biol. Cell 1–9 (2022). 10.1111/boc.202200007.

9. S. Ovejero, et al., A sterol-PI(4)P exchanger modulates the Tel1/ATM axis of the DNA damage response. EMBO J. 42 (2023).

10. H. C. Dippold, et al., GOLPH3 Bridges Phosphatidylinositol-4-Phosphate and Actomyosin to Stretch and Shape the Golgi to Promote Budding. Cell 139, 337–351 (2009).

11. S. E. Farber-Katz, et al., DNA Damage Triggers Golgi Dispersal via DNA-PK and GOLPH3. Cell 156, 413–427 (2014).

12. A. Traven, J. Heierhorst, SQ/TQ cluster domains: concentrated ATM/ATR kinase phosphorylation site regions in DNA-damage-response proteins. BioEssays 27, 397–407 (2005).

13. B. Bishé, G. H. Syed, S. J. Field, A. Siddiqui, Role of Phosphatidylinositol 4-Phosphate (PI4P) and Its Binding Protein GOLPH3 in Hepatitis C Virus Secretion. Journal of Biological Chemistry 287, 27637–27647 (2012).

14. M. M. Ng, H. C. Dippold, M. D. Buschman, C. J. Noakes, S. J. Field, GOLPH3L antagonizes GOLPH3 to determine Golgi morphology. Mol. Biol. Cell 24, 796–808 (2013).

15. R. Rizzo, et al., Golgi maturation-dependent glycoenzyme recycling controls glycosphingolipid biosynthesis and cell growth via GOLPH3. EMBO J. 40 (2021).

16. J. van Galen, et al., Sphingomyelin homeostasis is required to form functional enzymatic domains at the trans-Golgi network. Journal of Cell Biology 206, 609–618 (2014).

17. P. Lujan, et al., Sorting of secretory proteins at the trans-Golgi network by human TGN46. Elife 12 (2024).

18. Y. Wakana, et al., A new class of carriers that transport selective cargo from the trans Golgi network to the cell surface. EMBO J. 31, 3976–3990 (2012).

19. L. Anandi, V. Chakravarty, K. A. Ashiq, S. Bodakuntla, M. Lahiri, DNA-dependent protein kinase plays a central role in transformation of breast epithelial cells following alkylation damage. J. Cell Sci. 130, 3749–3763 (2017).

20. B. P. C. Chen, M. Li, A. Asaithamby, New insights into the roles of ATM and DNA-PKcs in the cellular response to oxidative stress. Cancer Lett. 327, 103–110 (2012).

21. S. Wang, et al., A Role of Rab29 in the Integrity of the Trans-Golgi Network and Retrograde Trafficking of Mannose-6-Phosphate Receptor. PLoS One 9, e96242 (2014).

22. K. Chabin-Brion, et al., The Golgi Complex Is a Microtubule-organizing Organelle. Mol. Biol. Cell 12, 2047–2060 (2001).

23. J. Wu, et al., Molecular Pathway of Microtubule Organization at the Golgi Apparatus. Dev. Cell 39, 44–60 (2016).

24. A. Efimov, et al., Asymmetric CLASP-Dependent Nucleation of Noncentrosomal Microtubules at the trans-Golgi Network. Dev. Cell 12, 917–930 (2007).

25. A. Venkataravi, M. Lahiri, DNA damage leads to microtubule stabilisation through an increase in Golgi-derived microtubules. bioRxiv (2022). 10.1101/2022.08.29.505705.

26. G. Bastianello, et al., Cell stretching activates an ATM mechano-transduction pathway that remodels cytoskeleton and chromatin. Cell Rep. 42, 113555 (2023).

27. Y. Ichijima, et al., Phosphorylation of histone H2AX at M phase in human cells without DNA damage response. Biochem. Biophys. Res. Commun. (2005). 10.1016/j.bbrc.2005.08.164.

28. K. J. McManus, M. J. Hendzel, ATM-dependent DNA Damage-independent Mitotic Phosphorylation of H2AX in Normally Growing Mammalian Cells. Mol. Biol. Cell 16, 5013–5025 (2005).

29. G. Nelson, M. Buhmann, T. von Zglinicki, DNA damage foci in mitosis are devoid of 53BP1. Cell Cycle 8, 3379–3383 (2009).

30. R. C. Burgess, B. Burman, M. J. Kruhlak, T. Misteli, Activation of DNA Damage Response Signaling by Condensed Chromatin. Cell Rep. 9, 1703–17 (2014).

31. C. R. Hunt, et al., Hyperthermia activates a subset of ataxia-telangiectasia mutated effectors independent of DNA strand breaks and heat shock protein 70 status. Cancer Res. (2007). 10.1158/0008-5472.CAN-06-4328.

32. Z. Bencokova, et al., ATM Activation and Signaling under Hypoxic Conditions. Mol. Cell. Biol. (2009). 10.1128/mcb.01301-08.

33. J.-H. Lee, T. T. Paull, Direct Activation of the ATM Protein Kinase by the Mre11/Rad50/Nbs1 Complex. Science (1979). 304, 93–96 (2004).

34. J. H. Lee, T. T. Paull, ATM activation by DNA double-strand breaks through the Mre11-Rad50-Nbs1 complex. Science (1979). 308, 551–554 (2005).

35. Z. Guo, S. Kozlov, M. F. Lavin, M. D. Person, T. T. Paull, ATM Activation by Oxidative Stress. Science (1979). 330, 517–521 (2010).

36. J.-H. Lee, T. T. Paull, Cellular functions of the protein kinase ATM and their relevance to human disease. Nat. Rev. Mol. Cell Biol. 22, 796–814 (2021).

37. G. Galea, et al., Spatiotemporal dynamics of DNA repair proteins between the Golgi and nucleus maintain genomic stability. bioRxiv (2025). 10.1101/2022.10.17.512236.

38. Á. dos Santos, et al., DNA damage alters nuclear mechanics through chromatin reorganization. Nucleic Acids Res. 49, 340–353 (2021).

39. N. Kleckner, et al., A mechanical basis for chromosome function. Proceedings of the National Academy of Sciences 101, 12592–12597 (2004).

40. J. Perry, N. Kleckner, The ATRs, ATMs, and TORs Are Giant HEAT Repeat Proteins. Cell 112, 151–155 (2003).

41. S. Ovejero, C. Soulet, M. Moriel-Carretero, The alkylating agent methyl methanesulfonate triggers lipid alterations at the inner nuclear membrane that are independent from its dna-damaging ability. Int. J. Mol. Sci. 22 (2021).

42. T. Stiff, et al., ATM and DNA-PK Function Redundantly to Phosphorylate H2AX after Exposure to Ionizing Radiation. Cancer Res. 64, 2390–2396 (2004).

